# Longitudinal transcriptome-wide gene expression analysis of sleep deprivation treatment shows involvement of circadian genes and immune pathways

**DOI:** 10.1101/628172

**Authors:** Jerome C. Foo, Nina Trautmann, Carsten Sticht, Jens Treutlein, Josef Frank, Fabian Streit, Stephanie H. Witt, Carolina De La Torre, Steffen Conrad von Heydendorff, Lea Sirignano, Junfang Chen, Bertram Müller-Myhsok, Andreas Meyer-Lindenberg, Christian C. Witt, Maria Gilles, Michael Deuschle, Marcella Rietschel

## Abstract

**Background:** Therapeutic sleep deprivation (SD) rapidly induces robust, transient antidepressant effects in a large proportion of major mood disorder patients suffering from a depressive episode, but underlying biological factors remain poorly understood. Research suggests that these patients may have altered circadian molecular genetic ‘clocks’ and that SD functions through ‘resetting’ dysregulated genes; additional factors may be involved, warranting further investigation. Leveraging advances in microarray technology enabling the transcriptome-wide assessment of gene expression, this study aimed to examine gene expression changes accompanying SD and recovery sleep in patients suffering from an episode of depression.

**Methods:** Patients (N=78) and controls (N=15) underwent SD, with blood taken at the same time of day before, after one night of SD and after recovery sleep. A transcriptome-wide gene-by-gene approach was used, with a targeted look also taken at circadian genes. Furthermore, gene set enrichment, and longitudinal gene set analyses including the time point after recovery sleep, were conducted.

**Results:** Circadian genes were significantly affected by SD, with patterns suggesting that molecular clocks of responders and non-responders, as well as patients and controls respond differently to chronobiologic stimuli. Notably, gene set analyses revealed a strong widespread effect of SD on pathways involved in immune function and inflammatory response, such as those involved in cytokine and especially in interleukin signalling. Longitudinal gene set analyses showed that in responders these pathways were upregulated after SD; in non-responders, little response was observed.

**Conclusions:** Our findings emphasize the close relationship between circadian, immune and sleep systems and their link to etiology of depression at the transcriptomic level.

## Introduction

Therapeutic sleep deprivation (SD) rapidly induces robust antidepressant effects in a large proportion of major mood disorder patients suffering from a depressive episode (1–5). The effects of the treatment are transient as relapse is usually observed after recovery sleep. The mechanisms through which SD exerts its antidepressant effects nevertheless offer important insights into the biological factors involved in depression and antidepressant response, and have been the focus of recent research (6–10). Work in humans (11–14)as well as animals (15–17) consistently documents the effects of mistimed or insufficient sleep and sleep deprivation on circadian gene expression (such as *CLOCK, ARNTL [BMAL1], PER1, PER2, PER3, etc*.,) as well as on genes involved in related biological processes such as inflammatory, immune and stress response (18, 19). A prominent hypothesis about the antidepressant mechanism underlying SD is that it restores circadian rhythmicity which is often dysregulated in depression, via resetting clock gene transcription (20, 21). The well-controlled nature and rapidity of response to SD treatment (22) renders it a promising context to investigate associated biological measures such as gene expression.

While no systematic investigation of gene expression changes in depressed patients undergoing SD treatment has been conducted to date, the study of gene expression in major depressive disorder (MDD) has raised the idea that genes associated with MDD are enriched for inflammation and immune response pathways, which may be linked to the sleep disturbances observed in depression (23–25). Circadian rhythms are found in the majority of physiological processes and the immune system is no different, with alterations of these rhythms leading to disturbed immune responses (26). The immune system and circadian clock circuitry crosstalk, with immune challenges and mediators, such as cytokines, also feeding back to affect circadian rhythms (27). Cytokines, including chemokines, interferons and interleukins, are integral to sleep homeostat regulation and can modulate behavioural and physiological functions (28).

We recently conducted a naturalistic study which aimed to examine clinical and genetic factors predicting response to SD (29). This was conducted in a sample of major mood disorder inpatients experiencing a depressive episode (n=78) and healthy controls (n=15). Briefly, 72% of patients responded to SD. Responders and non-responders did not differ in self/expert assessed symptom ratings or chronotype, but mood differed. Response was associated with lower age and later age at lifetime disease onset. Higher genetic burden of depression was observed in non-responders than healthy controls, with responders having intermediate risk scores.

The present study now aimed to examine gene expression changes accompanying SD and recovery sleep during a depressive episode using a longitudinal design, looking at changes in peripheral blood gene expression in the same sample. Gene expression changes occurring after SD and recovery sleep, as well as associated with response and patient-control status, were explored. In a hypothesis-driven approach, expression patterns of circadian genes were investigated. For a systematic search, transcriptome-wide gene-by-gene and gene set enrichment analyses were performed, while longitudinal gene set analysis (30) explored dynamics in expression trajectories at the functional gene set level.

## Methods and Materials

### Participants

The present sample has been described elsewhere (29). Seventy-eight inpatients (34 females; age mean ± standard deviation = 43.54 ± 14.80 years) presenting with a depressive episode (unipolar, n = 71; bipolar I, n = 6; and bipolar II, n = 1) and on stable medication for ≥ 5 days participated in this study. Depression was diagnosed according to ICD-10 criteria. Patients were recruited from consecutive admissions to the depression unit of the Department of Psychiatry and Psychotherapy of the Central Institute of Mental Health (CIMH), Mannheim, Germany. Prescribed medication included typical and atypical antidepressants, lithium, and adjunct therapies (anticonvulsants, antipsychotics and sleeping agents). Fifteen healthy controls (8 females; 40.53 ± 15.90 years) with no history of psychiatric/somatic disorders were recruited through an online advertisement on the CIMH website. The criteria for inclusion and exclusion were the same as the criteria for patients, except controls needed to lack psychiatric diseases, which was evaluated via Structured Clinical Interview for DSM-IV Axis II disorders (SCID-II) prior to SD. The investigation was carried out in accordance with the Declaration of Helsinki and approved by the CIMH ethics committee. All participants provided written informed consent following a detailed explanation of the study.

### Sleep Deprivation

On Day 1, participants gave informed consent and entered the study (see **Figure 1**). SD was conducted from Day 2–Day 3, in small groups of 1-5 participants under staff supervision. Participants were free to move around during the night of the SD protocol and were supervised by staff and occupied with activities such as games and walks to ensure wakefulness. Resting in bed was not allowed and SD was carried out on the ward under regular ward lighting conditions (no bright or dim lighting). Food intake was not restricted and some patients consumed snacks (i.e. bread) during SD. Participants underwent the same protocol irrespective of habitual sleep/wake timing; patients followed ward routines (i.e. lights on, lights out times) prior to inclusion to the study. Participants remained awake from ~0600hrs on Day 2 to 1800hrs on Day 3 (36 hours). On Day 3, participants underwent recovery sleep from 1800-0100hrs. Sleep phase advance was then carried out, shifting sleep one hour forward each day until the patient’s regular sleep pattern was reached. Controls participated alongside patients. Response was assessed by the senior clinical researcher using the Clinical Global Impression Scale for Global Improvement/Change in the afternoon on Day 3.

**Figure 1.**
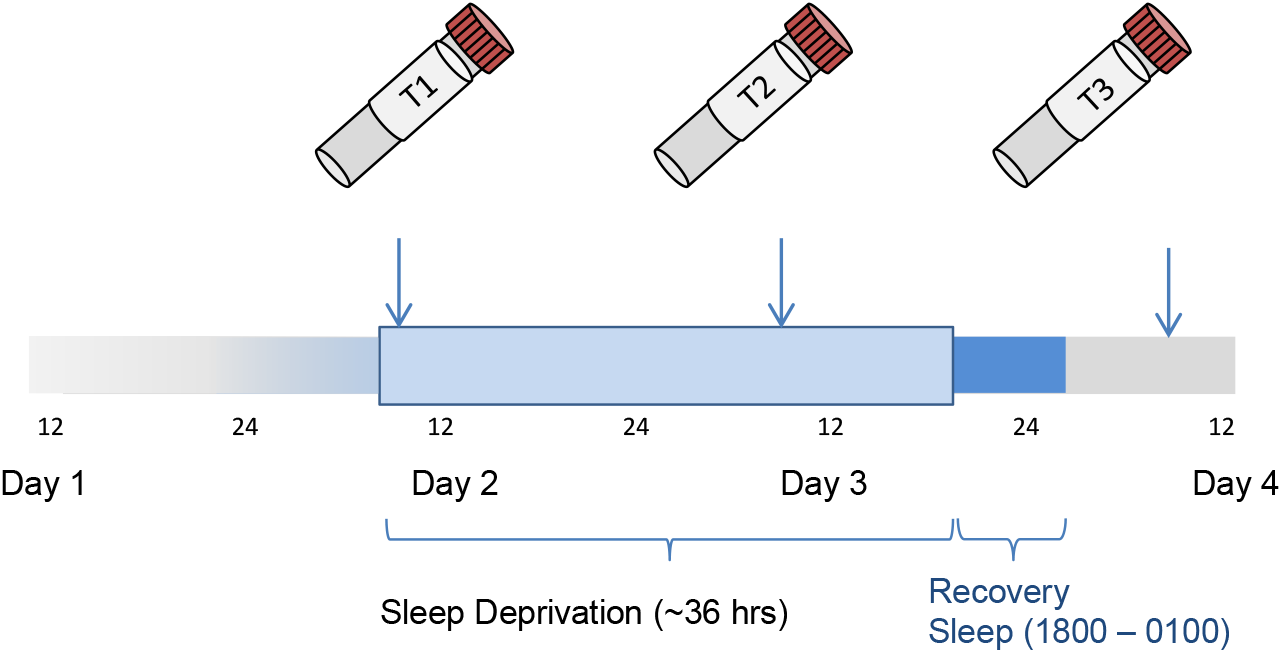
Experimental Procedure. Patients entered the study on Day 1 and underwent sleep deprivation for ~ 36 hours from Day 2 to Day 3 before undergoing recovery sleep. Response was assessed with the CGI-C in the afternoon of Day 3 before recovery sleep. Blood for gene expression was taken at the same time (0600-0730hrs) on Days 2, 3 and 4 (T1, T2, T3 respectively).

### Data Collection

Blood samples were collected in RNA-stabilizing PAXgene tubes (Qiagen, Hilden, Germany), processed according to standard procedures, and stored at −80□°C until analysis. Blood was collected at the same time (between 0600–0730hrs) on Day 2 before SD (T1), on Day 3 after SD (T2) and on Day 4 (T3) (see **Figure 1**). The number of samples decreased over time points (**Table 1**) due to nonparticipation.

**Table 1.** Samples included for analysis at different time points.

| Time Point | All Subjects | Patients | Responders | Non-Responders | Controls |
| --- | --- | --- | --- | --- | --- |
| T1 | 91 | 76 (43M/33F) | 60 (36M/24F) | 16 (7M/9F) | 15 (7M/8F) |
| T2 | 87 | 72 (41M/31F) | 56 (34M/22F) | 16 (7M/9F) | 15 (7M/8F) |
| T3 | 81 | 66 (38M/28F) | 53 (32M/21F) | 13 (6M/7F) | 15 (7M/8F) |
| Total | 259 | 214 (122M/92 F) | 169 (102M/67F) | 45 (20M/25F) | 45 (21M/24F) |

### Sample preparation and analysis

#### RNA and Microarray Analyses

Laboratory analyses were performed using standard methods (see **Supplementary Information, SI**).

#### Gene Expression Data Pre-processing

A Custom CDF Version 20 with ENTREZ based gene definitions was used to annotate the Affymetrix GeneChip™ Human Gene 2.0 ST arrays used for gene expression profiling (31). The Raw fluorescence intensity values were normalized applying quantile normalization and RMA background correction using SAS JMP11 Genomics, version 7(SAS Institute, Cary, NC, USA). The final dataset comprised mRNA expression targeting 24,733 unique genes for each time point per participant.

### Data Analysis

Analyses were conducted in R (Microsoft R Open 3.4.2). Significance was set at FDR q < 0.05.

#### Gene-based Analysis

Analyses were conducted using lme4 (Version 1.1-17). Linear mixed effects models with random intercepts were fitted to examine gene expression differences between T1 and T2 (‘effect of time point’). Three main models were fitted: effect of time point in ‘all patients’ (M1), in ‘responders vs. non-responders’ (M2) and in ‘patients vs. controls’ (M3). In all analyses, age and sex were included as covariates.

Models were specified as follows: for each transcript, likelihood ratio tests were calculated between two models (h1 vs h0). In both models, gene expression was specified as the dependent variable, covariates were specified as fixed effects and the individual was specified as a random effect, hi contained the comparison of interest specified as a fixed effect while h0 was a reduced model without it. That is, in M1, the ‘effect of time point’ was the only difference between h1 and h0, while in M2 and M3, the effect of interest was the interaction of the comparison group status (i.e. responder/non-responder, and patient/control) and ‘effect of time point’.

Additionally, whether differences in expression levels at T1 ‘baseline’ were informative about response and disease status was examined using linear models, controlling for sex and age.

#### Targeted Examination of Differential Expression of Circadian Genes

To determine differential expression of circadian genes, a list of genes comprising both traditional clock genes (in order of highest signal-to-noise ratio predicting circadian rhythmicity): *PER1, NR1D2, PER3, NR1D1, PER2, ARNTL, NPAS2, CLOCK, CRY2*, and *CRY1* as well as other top genes shown to predict circadian rhythmicity in human blood *(DDIT4, CLEC4E, FKBP5, DAAM2, TPST1, IL13RA1, SMAP2, HNRNPDL, FOSL2, PER1, FLT3, CDC42EP2, TMEM88, NR1D2, RBM3*) in (32) was used to take a focused look at results of the gene-based analysis. Using a Monte-Carlo approach, the probability of obtaining at least the observed number of significant associations (p < 0.05 uncorrected) in a random gene set of the same size was calculated.

#### Gene Set Enrichment Analysis

Gene Set Enrichment Analysis (GSEA)(33) was used to determine whether differentially expressed genes offered biologically meaningful insights about SD. Ranked lists were created based on results of models M1-M3, using a signed log10 transformed p-value with sign denoting direction of change, as described elsewhere(34). To allow a concise interpretation of the potentially widespread effects of SD interventions, the Hallmark gene set collection (35) (MSigDB Version 6.2), comprising 50 gene sets representing specific well-defined biological states/processes displaying coherent expression, was used. The heme metabolism gene set was excluded due to the globin interference artefact (communication with ThermoFisher).

#### Longitudinal Gene Set Trajectory

Time-course Gene Set Analysis (TcGSA)(30) (Version 0.12.1) was employed to examine gene expression dynamics over all three time points. TcGSA, employing mixed models, detects gene sets in which expression changes over time, taking between-gene and individual variability into account, with higher sensitivity and better interpretability than univariate individual gene analysis (for details, see(30)). As above, the Hallmark gene set collection was used. Significance was set at FDR q < 0.05. TcGSA was employed separately for all patients, controls, responders and non-responders. Data at T3 from two patients not following the recovery sleep protocol were excluded, while 8 participants had dropped out.

## Results

### Gene-based Analysis

In all patients [M1], 4,071 (2,083 up, 1,988 down) genes were significantly differentially expressed after SD. Significant differential changes in gene expression after SD between responders and nonresponders [M2] were observed in 360 genes [150 up, 210 down] and in patients vs. controls [M3] in 495 genes [248 up, 247 down].

**Table 2** shows top differentially expressed genes for these models. **<u>Tables S1.1-S1.3</u>** show the number of genes differentially expressed, upregulated, and downregulated for all models, and detailed lists of differentially expressed genes.

**Table 2.** Top 10 Differentially Expressed Genes for Each Model (T1 vs. T2)

| **M1 Effect of Time Point in Patients** |  |  |
| --- | --- | --- |
| Gene Symbol | Estimate | P-val FDR |
| TSPAN2 | 0,554523443 | 7,53143E-19 |
| KLF6 | 0,162981029 | 2,03907E-16 |
| MAK | 0,476984074 | 1,20412E-15 |
| ANTXR2 | 0,199530713 | 4,42924E-15 |
| TMEM43 | 0,19700767 | 5,80393E-15 |
| TREM1 | 0,300131404 | 1,1734E-14 |
| LRRFIP1 | 0,178631499 | 1,1734E-14 |
| NHSL2 | 0,253418357 | 2,50255E-14 |
| ARHGEF40 | 0,32560393 | 3,23301E-14 |
| SIPA1L1 | 0,228865158 | 3,2956E-14 |

| **M2 Response Status x Time Point** |  |  |
| --- | --- | --- |
| Gene Symbol | Estimate | P-val FDR |
| DNER | 0,19414884 | 0,000170023 |
| LPCAT2 | 0,32653286 | 0,000170023 |
| SLC10A5 | -0,62752802 | 0,000170023 |
| PCID2 | -0,300504 | 0,000209042 |
| TESC-AS1 | -0,3069843 | 0,001180095 |
| BECN1 | 0,21458155 | 0,001180095 |
| IFT74 | -0,43960958 | 0,001375265 |
| ZNF790 | -0,29221374 | 0,002668032 |
| GSR | 0,17871771 | 0,00297074 |
| LINC01125 | -0,16389193 | 0,00297074 |

| **M3 Patient-Control Status x Time Point** |  |  |
| --- | --- | --- |
| Gene Symbol | Estimate | P-val FDR |
| ERN1 | 0,30163938 | 0,003013102 |
| EZH2 | 0,38906547 | 0,003013102 |
| SLC44A1 | 0,27499289 | 0,003013102 |
| MEMO1 | 0,43613957 | 0,003013102 |
| UTP11L | 0,4553397 | 0,003013102 |
| NSFL1C | 0,2298895 | 0,003013102 |
| PCTP | 0,30731503 | 0,003013102 |
| ZBTB16 | 0,53874494 | 0,003013102 |
| CISH | -0,31446529 | 0,003456021 |
| TRAV4 | -0,4598706 | 0,003456021 |

FDR = false discovery rate

At baseline, no genes were significantly differentially expressed between patients/controls and responders/non-responders (**<u>Table S1.4</u>**).

### Target Examination of Differential Expression of Circadian Genes

We observed significant differential expression of circadian genes in models M1, M2 and M3. (**Table 3**, for more details, see **<u>Tables S2.1-3</u>**). Baseline differences in circadian genes between responders/non-responders, and patients/controls did not reach FDR q < 0.05, but achieved nominal significance (see **<u>Table S2.4</u>**).

**Table 3.** Differential Expression in Circadian Genes after SD

|  |  | Patients [M1] | Responders vs. Non-responders [M2] | Patients vs. Controls [M3] |
| --- | --- | --- | --- | --- |
|  | *Gene* | T1 vs. T2 | T1 vs. T2 | T1 vs. T2 |
| | | P-val($\beta$) | P-val($\beta$) | P-val($\beta$) |
| Traditional Clock Genes | PER1* | NS | **0.083 FDR (0.27)** | **0.045 FDR (0.32)** |
|  | NR1D2* | NS | NS | NS |
|  | PER3 | **0.0063 FDR (-0.11)** | NS | **0.065 FDR (-0.22)** |
|  | NR1D1 | **0.0014 FDR (-0.12)** | **0.035 unc (-0.14)** | **0.036 FDR (-0.27)** |
|  | PER2 | **0.038 unc (-0.059)** | NS | NS |
|  | ARNTL | **0.017 FDR (0.059)** | **0.033 unc (0.096)** | NS |
|  | NPAS2 | NS | NS | **0.0054 unc (-0.18)** |
|  | CRY2 | NS | NS | NS |
|  | CRY1 | NS | NS | NS |
|  | CLOCK | NS | **0.029 unc (-0.15)** | NS |
| Circadian Genes | DDIT4 | **1.24e-04 FDR (-0.32)** | NS | NS |
|  | CLEC4E | **0.0023 FDR (0.20)** | **0.0483 FDR (0.42)** | **0.018 FDR (0.48)** |
|  | FKBP5 | **0.037 unc (0.088)** | NS | **0.028 FDR (0.38)** |
|  | DAAM2 | **0.037 FDR (0.069)** | **0.020 unc (0.14)** | NS |
|  | TPST1 | **1.85e-04 FDR (0.23)** | NS | **0.025 FDR (0.44)** |
|  | IL13RA1 | NS | **0.0047 unc (0.19)** | NS |
|  | SMAP2 | **1.16e-5 FDR (0.12)** | **0.0390 FDR (0.18)** | **0.030 FDR (0.18)** |
|  | HNRNPDL | **1.32e-04 FDR (-0.10)** | **0.0173 unc (-0.13)** | **0.035 FDR (-0.18)** |
|  | FOSL2 | **2.23e-06 FDR (0.19)** | **0.0046 unc (0.22)** | **0.0046 unc (0.22)** |
|  | FLT3 | NS | NS | NS |
|  | CDC42EP2 | **4.19e-04 FDR (0.18)** | **0.0071 unc (0.26)** | NS |
|  | TMEM88 | **9.50e-05 FDR (0.21)** | NS | **0.023 unc (0.23)** |
|  | RBM3 | **7.12e-11 FDR (-0.21)** | NS | **0.024 unc (-0.14)** |

\*PER1 and NR1D2 are also on the list of circadian genes identified in Hughey et al. 2017.
FDR = false discovery rate, NS = not significant; unc = uncorrected
Blue = downregulation, Red = upregulation
**Bold** text indicates FDR $q < 0.05$

Monte Carlo simulations confirmed that significantly more circadian genes were differentially expressed than random gene sets of the same size (see <u>**SI**</u>).

### Gene Set Enrichment Analysis

GSEA notably found enrichment in immune response related pathways (see **<u>Tables S3.1-3</u>**). For M1, M2 and M3, 12, 23 and 11 gene sets were significantly positively enriched, respectively (FDR q < 0.05). The *TNFα signalling via NFKβ* gene set had the strongest positive enrichment in all models, while *Inflammatory Response* was also consistently among the significantly positively enriched gene sets.

Given the enrichment observed in Tumor Necrosis Factor Alpha (TNF*α*) and immune pathways, selected genes prominent in immune processes(25) were further examined (**<u>Table S4</u>**).

### Time course Gene Set Analysis

TcGSA results mirrored and extended gene-based analysis results:

In responders, 48 gene sets varied significantly (the model for one gene set, G2M Checkpoint, did not converge) (see **Figure 2a**). Descriptively, in comparison to T1, responders showed a spectrum of differential gene expression at T2, with strong upregulation observed (TNFα Signalling via NFKβ, IL6-JAK-STAT3-Signaling, Inflammatory Response, Angiogenesis) and maintained until T3. The Interferon Gamma Response and Interferon Alpha Response gene sets showed the strongest upregulation at T3.

**Figure 2.**
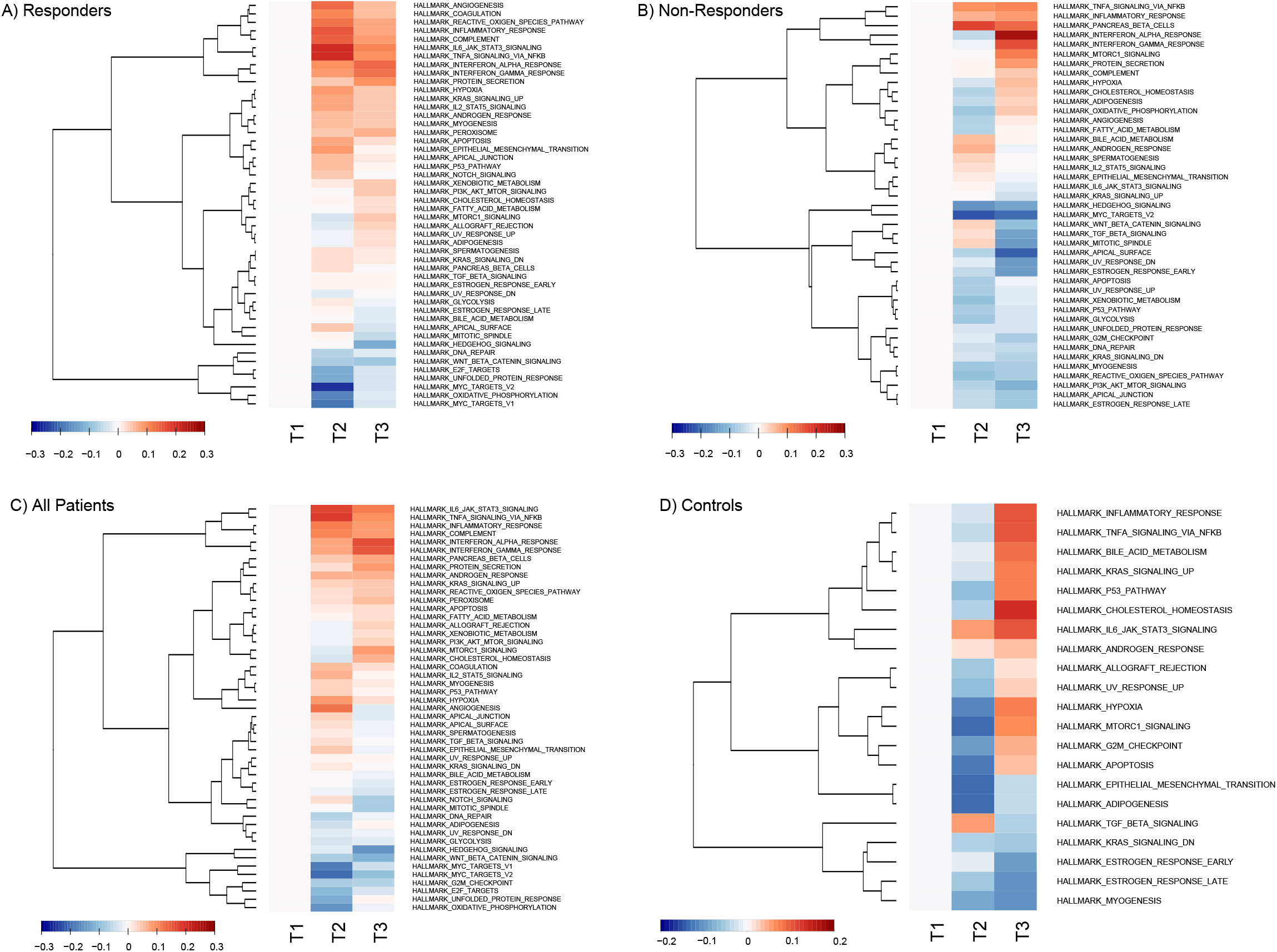
Heatmaps of estimated dynamics from significant gene sets in A) responders, B) nonresponders, C) patients, and D) controls. The median gene expression over subjects is used for each trend. Each trend is zeroed at T1 to represent baseline expression. Each row is a group of genes having the same trend inside a gene set, while the columns are time points. Trends are hierarchically clustered. Trends become red as median expression is upregulated or blue as it is downregulated compared to the baseline value at T1. The colour key represents the median of the standardized estimation of gene expression over the group of participants for a given trend in a significant gene set. Non-converging gene sets were excluded. For non-responders, the gene set for peroxisome was excluded from visualisation due to non-homogeneous expression within the set.

In non-responders, 44 gene sets varied significantly (the model for one gene set, Allograft Rejection, did not converge)(see **Figure 2b**), the majority of gene sets were downregulated at T2, with upregulation of the abovementioned gene sets only observed at T3.

In patients, expression was observed to vary significantly in 49 gene sets (see **Figure 2c**). Descriptively, upregulation was observed in immune system related gene sets in two main patterns; (1) strong upregulation at T2, sustained but weakening at T3 (i.e. Inflammatory Response, IL6-Jak-Stat3-Signalling and TNFα Signalling via NFKβ) and (2) light upregulation at T2 followed by stronger upregulation at T3 (e.g., Interferon Gamma Response, Interferon Alpha Response).

In controls, 21 gene sets varied significantly (**see Figure 2d**). Descriptively, at T3, immune/inflammation related gene sets were upregulated (e.g., Inflammatory Response, IL6-Jak-Stat3-Signalling, TNFα Signalling via NFKβ). The majority of other gene sets were downregulated at T2, followed by upregulation at T3.

**<u>Tables S5.1-4</u>** list significant TcGSA gene sets.

## Discussion

Here, we report the first longitudinal transcriptome-wide study of SD treatment in major mood disorder patients suffering from a major depressive episode. Widespread differential gene expression was observed after SD; circadian genes were differentially expressed, and enrichment in pathways related to immune function, inflammatory response and sleep regulation was observed.

It has been hypothesized that SD exerts its antidepressant effect by resetting disturbed clock gene functioning thought to be a feature of depression (9). Consequently, we examined changes in expression of circadian genes. All individuals, i.e. patients and controls, showed significant differential expression of circadian genes after SD. In responders, many significant gene expression changes were observed while in non-responders only a few nominally significant changes were found. The following observation may be of interest: three circadian genes, *PER1, CLEC4E* and *SMAP2* (FDR q = 0.083 approaching significance, 0.048, and 0.039, respectively) showed strong differential gene expression, i.e. increased expression, in responders versus non-responders after SD. While the roles of *CLEC4E* and *SMAP2* are yet unexplored in this context (however, see below for additional discussion of *CLEC4E*), similar *PER1*-related findings have been previously reported; consistent with the present findings, *PER1* expression is observed to be increased in SD responders and decreased in non-responders (9). Animal studies have shown that sleep deprivation or prolonged wakefulness enhanced *Per1* expression in several brain regions (20, 36), and that quetiapine increased *Per1* expression in the amygdala (37).

The present results are interesting in light of a study in human post-mortem brain tissue of ~12,000 ranked genes according to robustness of circadian rhythmicity across 6 brain regions; the top ranked circadian genes in that study were *ARNTL, PER2, PER3* and *NR1D1* and dysregulation in MDD vs controls were observed in these genes (21). The present study observed that these same genes were the most affected (of the traditional clock genes) as a result of SD, supporting the idea that SD may act on the dysregulation of these genes in depression.

The present findings cannot yet demonstrate that clock gene dysregulation/normalization is at the core of the SD mechanism. However, they suggest that (genes of) the molecular clocks of responders/non-responders, as well as patients/controls respond differently to chronobiologic stimuli, which may be associated with treatment outcomes.

Among the top 10 genes most significantly differentially expressed after SD in patients [M1] were: i) *KLF6*, observed to be upregulated in individuals after experimental restriction in a genome-wide association study of sleep duration (38); (ii) *SIPA1L1*, expression of which was found to be increased in a study examining changes in military personnel at baseline and after improvement of sleep (39); (iii) *NHSL2*, the function of which is unknown, but which is located in a genomic region found to be differentially hydroxymethylated in a study of sleep deprivation in mice(40); (iv) *TREM1*, which has immune function(11); and (v) *TSPAN2*, of which a study looking for gene expression changes under fluoxetine in rats found hippocampal upregulation (41). Among the top 10 genes observed in responders vs non-responders [M2] is *BECN1* (Beclin1) – which (together with *FKBP5*, see below) is shown to be involved in priming autophagic pathways to set the stage for antidepressant action (42, 43). Circadian rhythms of autophagic proteins, including beclin1 have been linked to sleep disturbances (44). Also in the top 10 in M2 were *DNER*, and *GSR* which have functions related to immune response(45, 46). Between patients and controls [M3], top genes included *EZH2*, which is reported to have a close relationship to IL-6 (47). *ZBTB16* is implicated in human sleep duration(38) and sleep deprivation in animal studies (48), and *CISH* is shown to be involved in immune related processes (49, 50). Among other differentially expressed genes are ones evidently associated with antidepressive intervention, such as *FKBP5* (51) where we observed significant upregulation in patients vs. controls after SD (see **<u>Tables S1.1. S2.1</u>**). *FKBP5* plays an important regulatory role in stress response (51–54); circadian rhythm abnormalities have been linked to the stress response system, and circadian clock genes can both regulate and be regulated by rhythms of glucocorticoid release (55).

Pathway analyses showed a global effect of SD on gene expression; immunological, inflammatory response and sleep regulation involved pathways were most strongly affected. These findings are of interest in light of previous gene expression studies linking immune function to MDD (56, 57); SD may affect pathways involved in the MDD etiology. In patients and especially responders, in the *Inflammatory Response* (genes related to cytokines, growth factors, cell differentiation markers and transcription factors)(35), *IL6-JAK-STAT3-Signalling* (aberrantly hyperactivated in patients with cancer and chronic inflammatory conditions)(58), and *TNFa Signalling via NFKβ* (cell proliferation, differentiation, apoptosis, neuroinflammation mediated cell death) gene sets, strong upregulation was observed at T2. Prior findings have shown associations between the immune system and depression, suggesting that causal pathways exist from immune dysregulation and inflammation to MDD (23, 24, 59–62). This raises the question of how SD and the immune system interact and whether SD counteracts and/or enhances depression-related immunological processes.

Interestingly, and in contrast to responders, mainly weaker downregulation was observed at T2 in non-responders, with upregulation of immune related gene sets only observed at T3. This differential function of immune related processes (i.e. blunted system responsiveness) might be associated with non-response to treatment. This preliminary finding must be further explored in larger sample sizes.

Of note, downregulation of *MYC targets* pathways after SD was consistently observed in both responders and non-responders but not controls. *MYC*, well known as an oncogene (63), acts as a mediator and coordinator of cell behaviour which also inversely modulates the impact of the cell cycle and circadian clock on gene expression (64). It has been shown that dysregulation of *MYC* disrupts the molecular clock by inducing dampened expression and oscillation of CLOCK-BMAL1 master circadian transcription factor (65). Removal of clock repression by *MYC* may play a role in the interaction of SD with depression and further investigation of these potential mechanisms is warranted.

Sleep and immunity are connected by anatomical and physiological bases (66). The role of cytokines in the brain is complex and remains to be fully understood; while a full discussion of their roles is beyond the scope of this work, **<u>Table S4</u>** shows cytokines and substances which have been implicated in both immune and sleep regulatory processes (25). The present findings, observing upregulation after SD, support reports linking levels of pro-inflammatory cytokines to depression and suggesting involvement in disease pathogenesis (25, 67, 68).

One important clue to SD response may lie in the observed TNFα expression patterns. TNFα is a pro-inflammatory cytokine controlling expression of inflammatory genetic networks; in addition to many immune-related functions in the brain, it influences whole organism function, including sleep regulation (25, 60). After SD, patients had significantly decreased expression of *TNF* compared to controls (FDR q =0.01); upregulation was observed in controls and downregulation in patients. It should be noted here that the sample sizes were unbalanced, potentially introducing bias in the result, however, upregulated *TNF* in controls is consistent with reports of sleep deprivation-induced increases in TNFα levels in healthy people (69). The decrease observed in patients, and especially responders, may inform the mechanism of SD response-increased TNFα concentrations are reported to be a marker of depression and TNFα administration is reported to induce depressive symptoms (70).

Sleep-wake cycles are accompanied by changes in circulating immune cell numbers and disturbing the circadian cycle affects immune response (71, 72). Acute SD reportedly affects diurnal rhythmicity of cells, mirroring the immediate immune stress response (73, 74). Consistent with a stress-like response, upregulation of expression of *IL6* and *IL1B*, thought to be involved in depression (75) and regulation of circadian rhythm/sleep homeostasis (12, 68, 76), was observed after SD. Given the tight coupling between circadian and immune systems, it may be that like the ‘resetting’ of circadian genes, the altered rhythmicity of the stress immune response system in depression is transiently normalized, leading to antidepressant effects; chronotherapeutic approaches that extend SD effects may be prolonging this normalization (20).

In support of the idea of involvement of stress response associated with SD is the fact that several of the differentially expressed genes are known to be driven by glucocorticoid signalling. One possibility is that differential expression observed has been induced by stress responses associated with SD; it is not possible in the present protocol to disentangle stress and SD. For example, while as indicated above *CLEC4E* is not explored in the SD context, it is known to be involved in immune function and also to be driven by glucocorticoids and is thus related to the stress response (77). Animal studies have shown that by removing glucocorticoid signalling via adrenalectomy, PER1, for example, was no-longer affected by SD, and responds to the stress induced by SD rather than SD itself (78).

There are several limitations to our study. The sample sizes of non-responders and controls were limited, with less power potentially contributing to the lower number of significant genes. The differences in sample sizes might furthermore create bias in the analyses of differential gene expression. While the statistical approach (linear mixed models) used in the single-gene analysis and underlying TcGSA is robust against unbalancedness and missing data, caution should still be exercised when considering the comparative results, as mentioned above. Next, although blood was sampled once per day on three consecutive days, a significant but manageable load for patients, the amount of data is sparse for statistical estimation. To better leverage longitudinal data and methods, a denser sampling scheme (e.g. every few hours) will be required to attain a more refined understanding of underlying SD mechanisms. It should be noted that the recovery sleep episode was a total of 7 hrs in length (i.e. 1800 to 0100 hrs) after a 36 hr homeostatic buildup of sleep pressure; this may not be long enough for homeostasis to recover and thus SD could have still had an effect on gene expression at T3. To address this in more detail, however, a different experimental design would have been required. Also, while food intake was not given special emphasis in this study, food intake can be an entrainment signal for the circadian clock. In the present study, some participants had small or simple meals (i.e. bread) and food intake was not restricted; possible effects cannot be excluded in the present design. Another direction which would be informative about circadian phase would be to include a marker of the central clock phase, e.g. melatonin secretion, also sampled at high temporal resolution. The sample studied was a naturalistic sample recruited from consecutive admissions and treated following standardized clinical guidelines. Medication regimens were tailored to the individual based on specific need, resulting in a variety of therapies used. While the effects of particular medications were not examined, to control for effects on results of the study, as an inclusion criterion, it was stipulated that the patient had to be on stable medication for at least 5 days prior to SD. Given the variety of medications used it was not possible to test for associations with drug response with sufficient statistical power. However, there was no apparent difference in substance class across response status. Nevertheless, robust effects of SD were observed, perhaps in part attributable to the consistent methodology for applying SD treatment in a relatively large cohort for SD. Considering the fact that depression is a heterogeneous phenotype accompanied by a heterogeneous immunophenotype (60), means that even larger sample sizes will be needed to substantiate the present findings. Finally, we investigated peripheral tissue transcriptome-wide expression changes; although an easily obtained, valuable proxy, the correlation of expression in the blood with expression in the brain, where depression is thought to act, is imperfect and requires further study (79). On the other hand, it is precisely these cells that best represent the current status of the immune system and the inflammatory response, the gene expression of which appears at the centre of SD. In addition, the crosstalk of the organs and biorhythm tuning is also mediated via the blood system.

Our findings affirm and emphasize the close relationship between circadian, immune and sleep systems in depression at the transcriptomic level, but the directionality of cause-effect remains unclear. Circadian, immune and sleep dysregulation may precede, accompany or come as a result of depression; they now represent targets for treatment which have the ability to influence clinical outcomes. Closer investigation of these systems with larger sample sizes and denser, longer-term sampling schemes will be key to disentangling and understanding the multi-level interactions occurring.

## Acknowledgements

This work has been supported by the Deutsche Forschungsgemeinschaft (DFG), SCHW 1768/1-1. AML acknowledges support from the German Federal Ministry of Education and Research (BMBF, grants 01ZX1314G, 01GS08147, 01EF1803A, 255156212 CRC 1158 subproject B09) and the Ministry of Science, Research and the Arts of the State of Baden-Wuerttemberg, Germany (MWK, grants 42-04HV.MED(16)/16/1, 42-04HV.MED(16)/27/1). MR was supported by the BMBF through grants within the e:Med research program 01ZX1314G. This work was also supported by BMBF/DLR grant 01EW1904 and BMBF / PTJ grant 031L0190A. This manuscript was uploaded to the preprint server bioRxiv.

