## Supplementary Info for "Longitudinal transcriptome-wide gene expression analysis of sleep deprivation treatment shows involvement of circadian genes and immune pathways"

**Supplementary Information:**

*Laboratory analyses*

RNA

RNA was isolated with the PAXgene Blood RNA Kit 50 (Qiagen, Hilden, Germany) according to standard protocols. Total RNA yield was determined using the Quant-iT RiboGreen RNA Reagent and Kit (Life Technologies, Darmstadt, Germany), and a Wallac Victor 2 1420 Multilabel Counter (Perkin Elmer, Rodgau, Germany). Total RNA purity was evaluated via optical density (OD) measurements (260 nm/280 nm) in a NanoDrop (peqLab, Erlangen, Germany). RNA integrity was determined by RNA integrity number measurement using RNA 6000 Nano Assay RNA chips run in an Agilent 2100 Bioanalyzer (Agilent Technologies, Santa Clara, CA, USA). The inclusion criteria were as follows: a ratio of 1.9–2.2 (OD 260/280); RNA integrity number>8.0, and the absence of a peak of genomic DNA contamination in electropherograms. Samples were transcribed to cDNA and hybridized to the Affymetrix GeneChip™ Human Gene 2.0 ST Array (Thermo Fisher Scientific Inc.) using the Whole Transcript Sense Target Labeling Assay protocol and 100 ng of total RNA from each sample.

*Microarray*

Gene expression profiling was performed using Affymetrix GeneChip™ Human Gene 2.0 ST arrays. Biotinylated antisense cRNA was prepared according to the Affymetrix standard labelling protocol with the GeneChip® WT Plus Reagent Kit and the GeneChip® Hybridization, Wash and Stain Kit. The chip was hybridized on an Applied Bioscience GeneChip Hybridization oven 640, dyed in the Applied Bioscience GeneChip Fluidics Station 450 and scanned with an Applied Bioscience GeneChip Scanner 3000 (all from Thermo Fisher Scientific Inc.).

Circadian Gene Set Monte Carlo Simulations

For effect of time point in all patients (M1), circadian genes were differentially expressed more frequently (p = 2.6 x 10^-4^ ) than a random gene set of the same size (Monte Carlo simulations: stabilising after 2,000,000 simulations) (**Figure S1a**). For effect of timepoint in responders vs. non-responders (M2), the circadian genes were differentially expressed more frequently (p = 9 x 10^-4^) than a random set of genes of the same size (250,000 simulations) (**Figure S1b**). For effects of time point in patients vs. controls, these genes were also differentially expressed more frequently (p = 1.2x10^-5^) than a random set of genes of the same size (1,500,000 simulations) (**Figure S1c**). Figures S1 plots cumulative probability during Monte Carlo simulations for Models M1-M3. **Table S3** shows significantly differential expression on circadian rhythmicity related genes for the different models.

**Figure S1.** Cumulative plot of Monte Carlo circadian gene overexpression test. a) effect of timepoint in all patients; b) effect of timepoint in responders vs. non-responders; and c) effect of time point in patients vs controls .

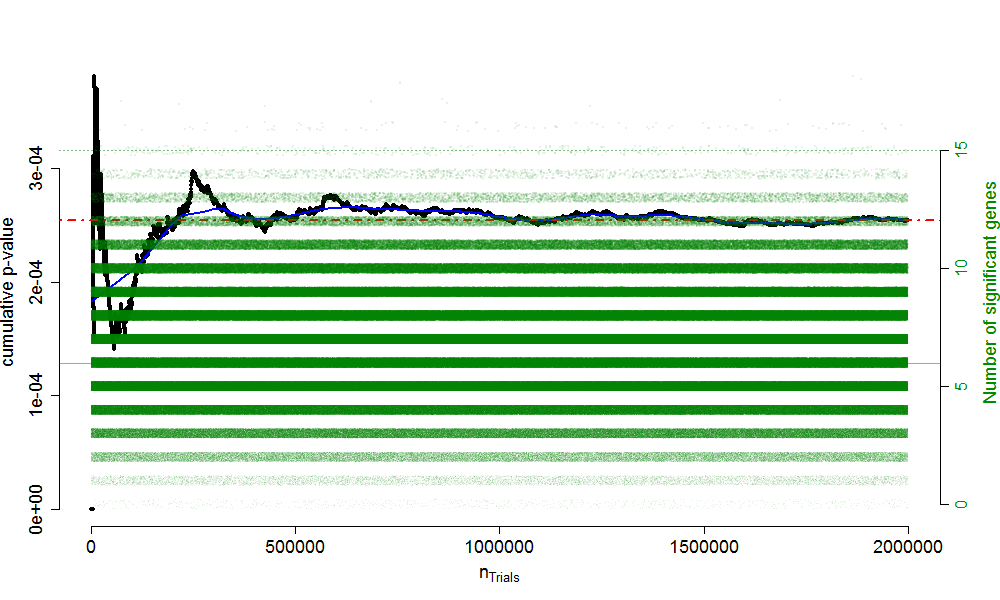

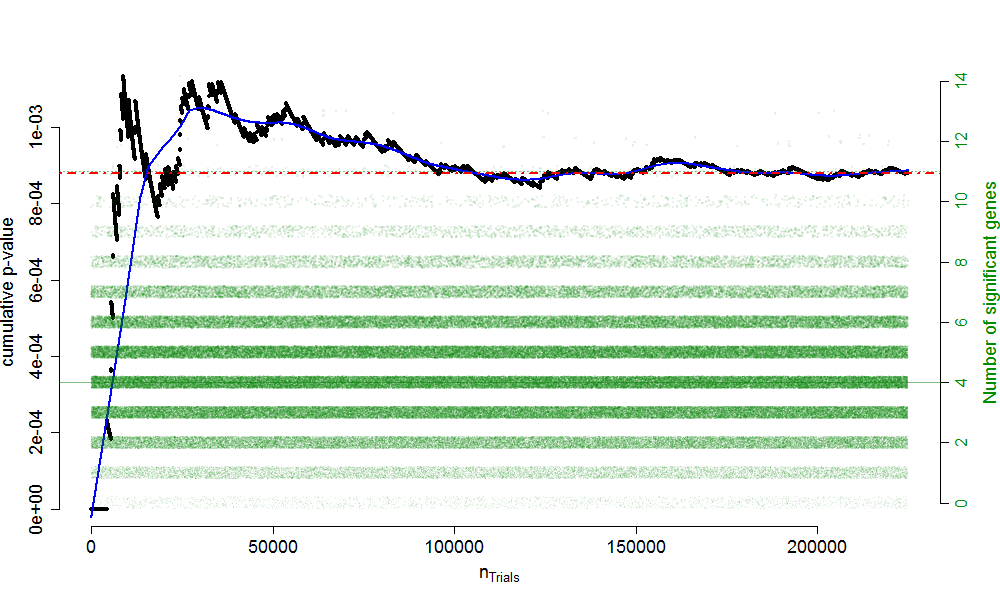

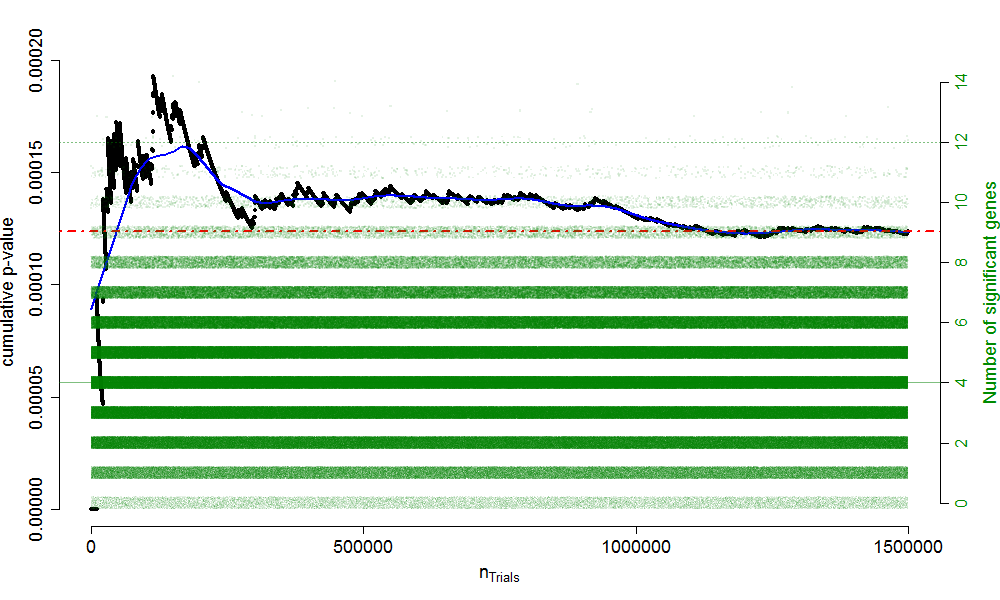
